# Electrophysiology of human iPSC-derived vascular smooth muscle cells and cell autonomous consequences of Cantu Syndrome mutations

**DOI:** 10.1101/2023.06.29.547088

**Authors:** Alex Hanson, Conor McClenaghan, Kuo-Chan Weng, Sarah Colijn, Amber N. Stratman, Carmen M. Halabi, Dorothy K. Grange, Jonathan R. Silva, Colin G. Nichols

**Affiliations:** Center for the Investigation of Membrane Excitability Diseases, Washington University in St. Louis, St. Louis, MO, 63130; Department of Cell Biology and Physiology, Washington University in St. Louis, St. Louis, MO, 63130; Department of Biomedical Engineering, Washington University in St. Louis, St. Louis, MO, 63130; Department of Pediatrics, Washington University in St. Louis, St. Louis, MO, 63130

**Keywords:** K_ATP_, KCNJ8, Kir6.1, ABCC9, SUR2, CRISPR, Cas9, cardiovascular system, whole cell patch clamp, I_Ca_, Cantu Syndrome, smooth muscle, VSMC, cardiac myocyte, blood pressure, vascular compliance, hiPSC, V_m_, LTCC, K_v_, BK, vascular tree, vascular tone, vasculopathy, cell-autonomous, RT-PCR, systemic vascular resistance

## Abstract

**Objective:** Cantu Syndrome (CS), a multisystem disease with a complex cardiovascular phenotype, is caused by GoF variants in the Kir6.1/SUR2 subunits of ATP-sensitive potassium (K_ATP_) channels, and is characterized by low systemic vascular resistance, as well as tortuous, dilated vessels, and decreased pulse-wave velocity. Thus, CS vascular dysfunction is multifactorial, with distinct hypomyotonic and hyperelastic components. To dissect whether such complexities arise cell-autonomously within vascular smooth muscle cells (VSMCs), or as secondary responses to the pathophysiological milieu, we assessed electrical properties and gene expression in human induced pluripotent stem cell-derived VSMCs (hiPSC-VSMCs), differentiated from control and CS patient-derived hiPSCs, and in native mouse control and CS VSMCs.

**Approach and Results:** Whole-cell voltage-clamp of isolated aortic and mesenteric VSMCs isolated from wild type (WT) and Kir6.1[V65M] (CS) mice revealed no difference in voltage-gated K^+^ (K_v_) or Ca^2+^ currents. K_v_ and Ca^2+^ currents were also not different between validated hiPSC-VSMCs differentiated from control and CS patient-derived hiPSCs. Pinacidil-sensitive K_ATP_ currents in control hiPSC-VSMCs were consistent with those in WT mouse VSMCs, and were considerably larger in CS hiPSC-VSMCs. Consistent with lack of any compensatory modulation of other currents, this resulted in membrane hyperpolarization, explaining the hypomyotonic basis of CS vasculopathy. Increased compliance and dilation in isolated CS mouse aortae, was associated with increased elastin mRNA expression. This was consistent with higher levels of elastin mRNA in CS hiPSC-VSMCs, suggesting that the hyperelastic component of CS vasculopathy is a cell-autonomous consequence of vascular K_ATP_ GoF.

**Conclusions:** The results show that hiPSC-VSMCs reiterate expression of the same major ion currents as primary VSMCs, validating the use of these cells to study vascular disease. The results further indicate that both the hypomyotonic and hyperelastic components of CS vasculopathy are cell-autonomous phenomena driven by K_ATP_ overactivity within VSMCs.

## Introduction

Cardiovascular events are often preceded by vascular abnormalities, and understanding fundamental vasculopathic mechanisms is critical to explaining disease progression. Interventional experimentation on *bona fide* VSMCs can realistically only be carried out in non-human animal models, but the potential for species-dependent differences to obscure human relevance ultimately warrants replication in human cell-based systems. VSMCs derived from human induced pluripotent stem cells (hiPSCs) are promising for the study of human-specific, as well as mutation-specific and cell-autonomous, mechanisms of cardiovascular disease, and for pharmacotherapeutic testing^1, 2^. The latter has generated excitement and is the subject of active, ongoing study in the field of hiPSC-derived 3D tissue engineering^3, 4^, but remains hampered by key obstacles, such as deficient *in vitro* elastin expression^5^. New approaches for differentiation of hiPSCs into VSMCs (hiPSC-VSMCs) are increasingly being used to shape our understanding of vascular biology and fundamental disease mechanisms^1, 2, 6^. The most robust differentiation protocols combine transcriptomic, metabolomic, proteomic, and bio-functional modalities to holistically characterize cell identity, and enable essentially identical reproducibility by merit of employing completely chemically defined conditions.

Notably, however, while the electrophysiology of cardiac myocytes derived from hiPSCs has been extensively studied^7^, hiPSC-VSMC electrophysiology (EP) has remained unexplored, leaving a critical aspect of hiPSC-VSMC function unknown. VSMC excitability directly determines myocyte tone in resistance vessels, and in turn systemic vascular resistance (SVR)^8^. Resistance vessel VSMCs are normally in a partially activated state that is dually responsive to myotonic stimulation or inhibition; small positive V_m_ deflections activate voltage-sensitive L-type Ca^2+^ (LTC) channels, while hyperpolarizing stimuli, such as activation of K^+^ channels, shift the V_m_ away from the activation range of LTC channels, resulting in reduced excitability and vasodilation. Elastic vessels, such as the aorta, accommodate cardiac stroke volume during systole, and passively recoil during diastole, delivering a non-pulsatile, laminar flow of blood to distal vessels. This passive function is conferred by the characteristic network of elastic fibers of large proximal vessels, which in turn is produced and deposited by elastogenic VSMCs. Increased elastin expression is a recognized consequence of pharmacological activation of vascular ATP-sensitive K^+^ (K_ATP_) channels^9–12^. Thus, electrical activity is a key component of normal VSMC function in resistance vessels as well as elastic vessels, and electrical abnormalities have been shown to dramatically alter VSMC function in ways that are relevant to disease ^13–16^. Therefore, it is critical to understand the EP of hiPSC-VSMCs, and to appreciate how it compares to, or may differ from, that of native VSMCs.

Vascular dysfunction underlies the complex cardiovascular phenotype of Cantú Syndrome (CS), which encompasses cardiac hypertrophy and hypercontractility, patent ductus arteriosus, low systemic blood pressure, aortic root dilatation, and decreased pulse-wave velocity^17–19^. CS is caused by gain-of-function (GoF) mutations in *KCNJ8* and *ABCC9*, which encode Kir6.1 and SUR2, respectively the pore and regulatory subunits of the vascular K_ATP_ channel: ^17, 20^. Importantly, cardiac features of CS are driven by K_ATP_ GoF within the vasculature, mediated by elevated renin-angiotensin signaling and baroreceptor-mediated adrenergic signaling^18, 21, 22^. CS vasculopathy is also characterized by striking abnormalities are also observed more proximally in the large, elastic vessels, which exhibit gross morphological changes such as dilatation and tortuosity^23, 24^, associated with abnormal biomechanical properties including excessive aortic compliance and decreased PWV^18, 20^. Thus, CS vasculopathy is multifactorial with distinct hypomyotonic and hyperelastogenic components, which together drive broader cardiovascular consequences via elevated renin-angiotensin and adrenergic signaling.

The present study was undertaken to address multiple issues raised above: (i) to address the current lack of electrophysiological characterization of hiPSC-derived VSMCs, and provide a first characterization of relevant vascular ionic currents; (ii) to assess any potential CS-dependent changes in other (i.e., non-K_ATP_) vascular currents, and (iii), to gain insight to the hypomyotonic and hyperelastic components of CS vasculopathy using native mouse VSMCs and hiPSC-VSMCs, and establish whether these phenomena arise as cell-autonomous consequences of VSMC K_ATP_ GoF, or secondarily to non-VSMC K_ATP_ GOF.

## Methods

### Study approval

Animal studies were performed in compliance with the standards for the care and use of animal subjects defined in the NIH Guide for the Care and Use of Laboratory Animals^26^ and were reviewed and approved by the Washington University Institutional Animal Care and Use Committee. Human studies were approved by the Washington University Human Studies Committee and carried out with the full consent of participating patients.

### Mouse lines

Generation of knock-in mice carrying human gain-of-function mutations in the *ABCC9* or *KCNJ8* genes using CRISPR/Cas9-mediated genome engineering technology were described previously ^18^. Briefly, guide RNA (gRNA) target sequences were predicted using the MIT CRISPR design tool (http://crispr.mit.edu). abcc9 gRNA (5’ CATTGCCACGAAGCTGGCGG 3’) or kcnj8 gRNA (5’ ACGCCACTTCAGGTCTACCA 3’) were cloned into BbsI digested plasmid pX330 (Addgene # 42230). T7 sgRNA template and T7 Cas9 template were prepared by PCR amplification and gel purification, and injected into zygotes from superovulated B6CBA F1/J female mice mated with B6CBA F1/J male mice. After injection, zygotes were incubated at 5.5% CO_2_ at 37 °C for 2 h, and surviving embryos were transferred to ICR recipient mice by oviduct transfer. We identified 12 positive founder animals carrying CS mutation mouse SUR2[A476V] (equivalent to human SUR2[A478V]), and two positive founder animals carrying the CS mutation Kir6.1[V65M]. All founders were viable and fertile. Successful mutation of founder (F0) mice was verified by Sanger sequencing of genomic DNA and mutant mice were subsequently crossed with C57BL/6J mice to generate heterozygous F1 Kir6.1^wt/VM^ and SUR2^wt/AV^ lines. PCR was used to generate amplicons of *KCNJ8* and *ABCC9* spanning >1 kb either side of the introduced mutation, from gDNA isolated from mouse tails, and resultant PCR products were sequenced to confirm absence of unintended additional mutations. After verification, one F1 animal from one line of each genotype was selected and further crossed against C57BL/6J to generate F2 Hets which were intercrossed to generate subsequent generation Het and Homo as well as wild type littermates that were used in experiments.

### Human iPSC generation and differentiation to VSMCs

CS patient-derived hiPSCs heterozygous for the RQ and RW variants were generated, respectively, from peripheral blood mononuclear cells and renal epithelial cells, which were generously provided by CS patients who had been genetically confirmed to carry these variants. Human R1154W patient renal epithelial cells (RECs) were reprogrammed to hiPSCs by the WUSM Genome Engineering and iPSC Core (GEiC) using Sendai Virus-based reprogramming vectors. After four unsuccessful attempts to reprogram human R1154Q patient RECs, peripheral blood mononuclear cells were provided by the patient, and were successfully reprogrammed by the GEiC using the Sendai Virus-based reprogramming cocktail. hiPSCs were maintained on a 4-day passaging cycle. Two subclonal hiPSC lines were generated for each patient sample, and DNA sequencing analysis confirmed the expected gene variant in each line. Two control lines, C2a (gift of Dr. Gordana Vunjak-Novakovic, Columbia University), and AN1.1 (gift of Amber Neilson) were used as controls. hiPSCs were differentiated to hiPSC-VSMCs by adapting the chemically-defined protocol published by Patsch et al.^77^, which involves initially ‘priming’ hiPSCs into lateral plate mesodermal progenitors through Wnt activation and hBMP4 treatment, with subsequent induction to VSMC fate though co-treatment with PDGF-BB and Activin A. The original protocol was followed exactly, except that differentiation was carried out in 12-well plates, rather than in T175 flasks. Cryopreserved human aortic smooth muscle cells were obtained from Lonza Pharma and Biotech (Catalog #: CC-2571). Cells were plated on gelatin-coated coverslips, and grown in DMEM with 10% FBS for 3-5 days before analysis. Immunohistochemical (IHC) staining for smooth muscle actin was carried out using anti alpha-Smooth Muscle-Cy3 monoclonal antibody C6198 (Sigma-Aldrich Co.).

### RNA analysis and immunocytochemistry in native VSMCs and hiPSC-VSMCs

For aortae and hiPSC-VSMCs, RNA was isolated using Trizol*®* following manufacturer’s protocol (Life Technologies, Grand Island, NY). For iPSCs, first strand cDNA was synthesized using SuperScript™ III First-Strand Synthesis System (Thermo Fisher). For aortic samples, 1 μg of total RNA was reverse transcribed using High-Capacity RNA-to-cDNA™ Kit, per manufacturer’s protocol (Life Technologies, Grand Island, NY). Quantitative real time PCR (qPCR) was done using 100 ng of cDNA template, TaqMan® Fast Universal PCR Master Mix and TaqMan assays (primers/probes) purchased from Life Technologies (Grand Island, NY). Reactions were run in duplicate on the QuantStudio-3 real-time PCR system (Applied Biosystems) using the Fast PCR parameters (95°C for 20 sec followed by 40 cycles of 95°C for 1sec and 60°C for 20 sec). Experimental gene expression was normalized to that of *Gapdh* or β2-microglobulin (*B2m*) transcript levels as indicated in the figures. Taqman assays used in this study are Mm00514670_m1 (*Eln*), Mm99999915_g1 (*Gapdh*) and Mm00437762_m1 (*B2m*).

For immunocytochemistry, cells were fixed with 4% PFA for 5 hours, permeabilized with 100% ice cold methanol for 10 min, blocked with 1% BSA, 22.52 mg/ml glycine in PBS with 0.1% Tween 20 (PBST) for 30 min and incubated in primary antibody diluted in 1% BSA/PBST for 1 hour. After washing with PBS, slides were incubated in secondary antibody with Hoechst 34580 (0.5 µg/ml; #63493; Sigma) in 1% BSA/PBST for 1 hr. After mounting with Fluoromount™ Aqueous Mounting Medium (F4680; Sigma), images were obtained using a 20x objective with a W1 Spinning Disk confocal microscope, a Fusion camera, and the Nikon Eclipse Ti2-E base. Images were then de-convolved using Nikon NIS-Elements software. Primary antibodies used for immunocytochemistry were: mouse anti-PDGFRβ (1:100; ab69506; Abcam), mouse anti-Smoothelin (1:100; sc-376902, AF647 conjugated; Santa Cruz), rabbit anti-MYH11 (1:100; ab82541; Abcam), rabbit anti-α-SMA (1:100; D4K9N, XP; Cell Signaling Technology), and rabbit anti-TAGLN (1:100; ab14106; Abcam). Secondary antibodies used were acquired from Invitrogen and used at a 1:750 dilution: Goat anti-Mouse IgG (H+L) Highly Cross-Adsorbed Secondary Antibody, Alexa Fluor™ 594 (A-11032), Goat anti-Rabbit IgG (H+L) Highly Cross-Adsorbed Secondary Antibody, Alexa Fluor™ 488 (A-11034), and Goat anti-Rabbit IgG (H+L) Highly Cross-Adsorbed Secondary Antibody, Alexa Fluor™ 633 (A-21071).

### Patch clamp electrophysiology

Mice were anesthetized with 2.5% avertin (10ml/kg, intraperitoneal Sigma-Aldrich) and the descending aorta or mesenteric arteries were rapidly dissected and placed in ice-cold Physiological Saline Solution (PSS) containing (in mM): NaCl 134, KCl 6, CaCl_2_ 2, MgCl_2_ 1, HEPES 10, and glucose 10, with pH adjusted to 7.4 with NaOH. Smooth muscle cells were enzymatically dissociated in dissociation solution containing (in mM): NaCl 55, sodium glutamate 80, KCl 5.6, MgCl_2_ 2, HEPES 10, and glucose 10, pH 7.3 with NaOH, then placed into dissociation solution containing papain 12.5 µg/mL, dithioerythreitol 1 mg/mL, and BSA 1 mg/mL for 25 minutes (at 37°C), before transfer to dissociation solution containing collagenase (type H:F=1:2) 1 mg/mL, and BSA 1 mg/mL for 5 minutes (at 37°C). Cells were dispersed by gentle trituration using a Pasteur pipette, plated onto glass coverslips on ice and allowed to adhere for >1 h before transferal to the recording chamber.

Whole-cell ion currents were recorded using an Axopatch 200B amplifier and Digidata 1200 (Molecular Devices). Recordings were sampled at 3 kHz and filtered at 1 KHz. Voltage-gated K currents were measured in High Na^+^ bath solution containing (in mM): NaCl 134, KCl 5.4, CaCl_2_ 100 μM, MgCl_2_ 1, HEPES 10, and glucose 10, with pH adjusted to 7.4 with NaOH. The pipette solution contained (in mM) KCl 140, MgCl_2_ 1, HEPES 10, glucose 10, and EGTA 10, ATP 5, with pH adjusted to 7.2 with KOH. Voltage-gated L-type Ca (LTC) currents were measured in High Na^+^ bath solution containing (in mM): choline chloride 124, BaCl_2_ 20, MgCl_2_ 1, HEPES 10, and glucose 5, with pH adjusted to 7.4 with NaOH. The pipette solution contained (in mM) CsCl 130, MgCl_2_ 2, HEPES 10, glucose 10, and EGTA 10, NaATP 3.5, with pH adjusted to 7.3 with KOH. To assess K_ATP_ conductances, currents were initially measured at a holding potential of -70mV in High Na^+^ bath solution containing (in mM): NaCl 136, KCl 6, CaCl_2_ 2, MgCl_2_ 1, HEPES 10, and glucose 10, with pH adjusted to 7.4 with NaOH before switching to a High K^+^ bath solution (KCl 140, CaCl_2_ 2, MgCl_2_ 1, HEPES 10, and glucose 10, with pH adjusted to 7.4 with KOH) in the absence and presence of pinacidil and glibenclamide as indicated. The pipette solution contained (in mM) potassium aspartate 110, KCl 30, NaCl 10, MgCl_2_ 1, HEPES 10, CaCl_2_ 0.5, K_2_HPO_4_ 4, and EGTA 5, with pH adjusted to 7.2 with KOH.

### Arterial compliance

After mice were euthanized under isoflurane anesthesia, the aortae of 3 week-old mice were excised and placed in a physiologic saline solution (PSS) containing (mM) 130 NaCl, 4.7 KCl, 1.18 MgSO_4_-, 1.17 KH_2_PO_4_, 14.8 NaHCO_3_, 5.5 dextrose, and 0.026 EDTA (pH 7.4). The vessels were then cleaned from surrounding fat, mounted on a pressure arteriograph (Danish Myo Technology) and maintained in PSS at 37°C. Vessels were visualized with an inverted microscope connected to a charged-coupled device camera and a computerized system, which allows continuous recording of vessel diameter. Intravascular pressure was increased from 0 to 175 mmHg by 25-mmHg increments and the vessel outer diameter was recorded at each step (12 seconds per step). The average of three measurements at each pressure was reported.

### Data Analysis

Unless otherwise noted, data are presented as mean ± S.E.M. The Real Statistics add-in package was used to run statistical analysis in Microsoft Excel. All data were tested for statistical significance using Mann Whitney *U* test, or Kruskal-Wallis test with post hoc Dunn’s test.

## Results

### Non-K_ATP_ ionic currents that govern vascular excitability are not altered in CS vasculopathy

VSMC excitability depends on the concerted activity of multiple ion conductances, in particular voltage-gated K^+^ (K_v_) currents and L-Type Ca^2+^ currents (LTCCs). To provide a benchmark of these conductances in native wild type VSMCs, whole-cell voltage-clamp was first used to assess functional activity of K_v_ channels in acutely dissociated WT mouse aortic VSMCs (Fig. 1). The voltage protocol was designed to inclusively measure the entire ensemble of VSMC K_v_ channels, with high (5mM) ATP included in the internal solution to exclude K_ATP_ currents. Very small, linear, K^+^ conductance was detected below ∼-25mV, but additional K^+^ conductance with apparently instantaneous and time-dependent components were increasingly activated at more positive voltages (Fig. 1A). Additionally, more positive voltage steps revealed observable single-channel activity consistent with large-conductance calcium-activated potassium (BK) channels (Fig.1A). To assess whether these vascular K_v_ currents are altered in CS vasculopathy, aortic VSMCs were then acutely dissociated from mice carrying the Kir6.1[V65M] (VM) mutation, a CS-associated variant that causes marked K_ATP_ GoF and severe CS vasculopathy^18, 27^. K_v_ currents in VM VSMCs were very similar to WT (Fig. 1), and summarized I-V curves show that overall K_v_ current amplitudes (measured during the final 100 msecs of the voltage pulse) were indistinguishable between WT and VM VSMCs (Fig. 1B). To separate K_v_ and BK currents, and to assess K_v_ current kinetics, a monoexponential function representing ‘idealized’ K_v_ current was fit directly to each recording at the three most positive voltage steps (i.e., +25mV, +35mV, and +45mV), as shown in Fig. 1C. This allowed separation of instantaneous K_v_ amplitudes, time-dependent K_v_ amplitudes, and assessment of K_v_ time constant (Tau). BK channel activity (NP_o_) was separately estimated for each record at +25mV, +35mV, and +45mV by subtracting the idealized K_v_ current from each raw recording. Instantaneous and time-dependent K_v_ amplitudes, as well as K_v_ kinetics, and BK activity, were all statistically unaltered in the setting of CS vasculopathy (Fig. 1C).

**Figure 1:**
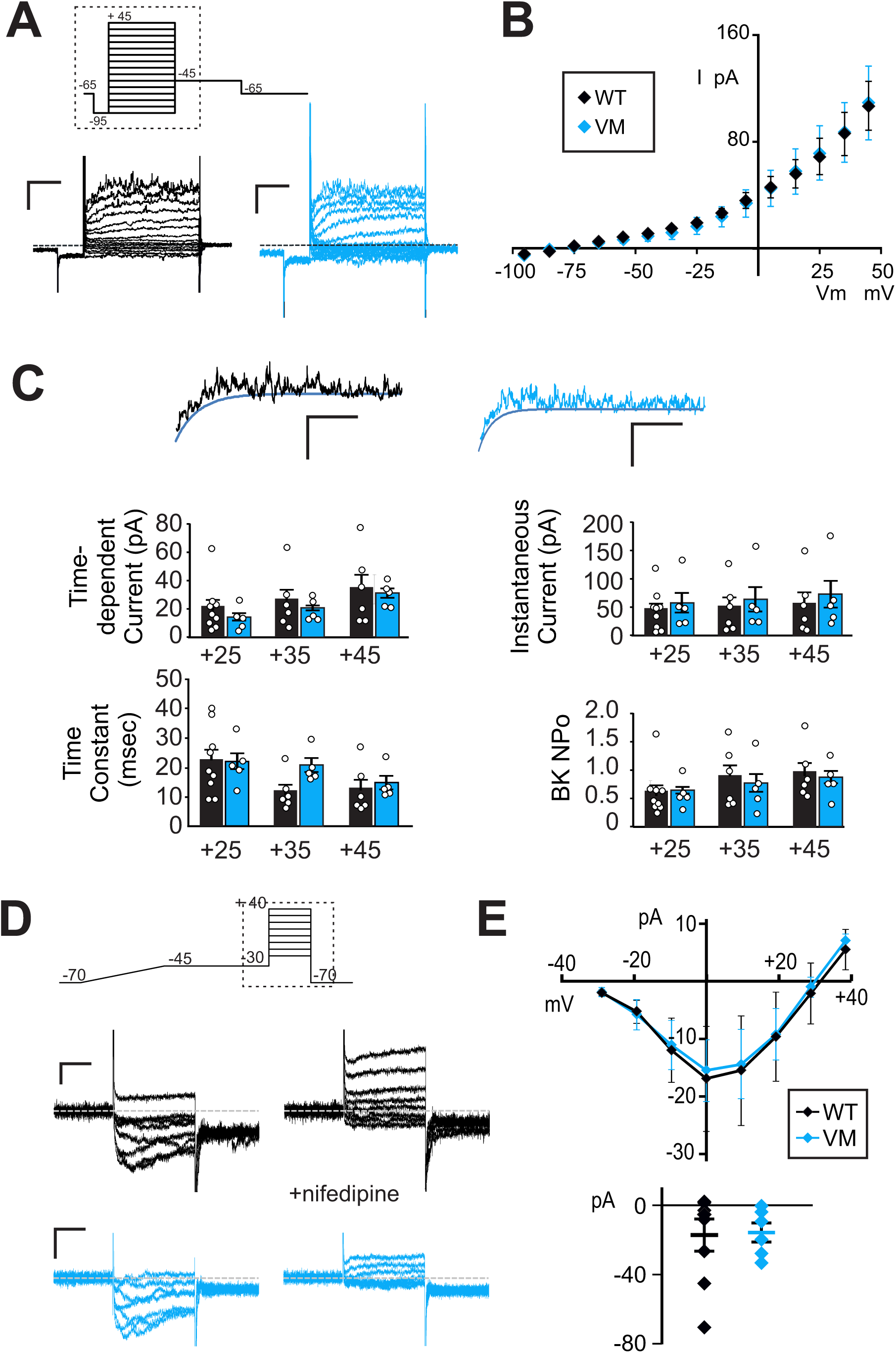
Voltage-gated K and Ca currents in mouse CS and WT VSMCs. (**A**) Representative whole-cell voltage-clamp recordings show K_v_ currents from acutely isolated aortic VSMCs from a WT (black) and VM (blue) mouse (scale bars = 50 pA, 100 ms). Voltage was stepped from holding potential of -65mV to pre-pulse at -95 mV, then stepped more positive in 10mV increments. Hashed box demarcates the portion depicted in the representative patch-clamp traces. (**B**) Peak I-V relationships (measured during the final 100 msecs of the voltage pulse) show mean ± standard error (s.e.m.), for n= 9 cells from four mice (WT) and 5 cells from three mice (VM) cells in each case. (**C**) Representative traces following step to +45 mV from WT (black) and VM (blue) VSMCs, fit with monoexponential functions to the minimal currents as a function of time, assumed to reflect K_v_ currents without the contaminating fluctuating, presumably BK current. From such traces, K_v_ amplitudes and kinetics, as well as BK currents, were calculated, as shown to the right (n as above). Statistical significance was determined by Mann-Whitney *U* test (α = 0.05, no significant differences detected). (**D**) Representative whole-cell voltage-clamp recordings of LTCCs were obtained from acutely isolated mesenteric VSMCs from WT (black) and VM (blue) mouse (scale bars = 10 pA, 75 ms). Voltage was ramped from - 70 mV to -45 mV over 400 msecs to inactivate any high voltage activated currents, then stepped to voltages between -30 and +40 mV in 10 mV steps. Hashed box demarcates the portion depicted in the representative patch-clamp traces. Recordings were obtained in the presence of 1μM Bay K8644 (*left*), and conductance by LTCC was confirmed by >95% inhibition with 10μM nifedipine (*right*). (**E**) (*above*) Peak I-V relationships on Bay K8644 are summarized, with (*below*) peak current at 0 mV for individual traces. Error bars show mean + s.e.m. (n = 8 cells cells from four WT mice and 6 cells from four VM mice (in each case). Statistical significance was determined by Mann-Whitney *U* test (α = 0.05, no significant differences detected).

Next, whole-cell voltage-clamp was used to assess functional activity of voltage-sensitive LTCC. Ba^2+^ was used as a charge carrier, and included in the bath was BayK8644 (1μM), a potentiator of vascular LTCCs. K^+^ currents were minimized by inclusion of Cs^+^ (130mM) in the internal solution, as well as 3.5mM ATP to exclude K_ATP_ currents. LTCCs were undetectable in acutely isolated WT mouse aortic VSMCs (n = 8 recordings from three animals). However, characteristic LTCCs that were fully inhibited by nifedipine (10μM) were present in VSMCs acutely dissociated from first-through fourth-order mesenteric arteries, which are resistance vessels (Fig. 1D). Again, LTCCs recorded from mesenteric VSMCs acutely dissociated from VM mice were essentially identical to WT (Fig. 1E).

Together, the above experiments indicate that, despite marked changes in VSMC K_ATP_ currents in Cantu mice^18^, there are no accompanying compensatory or downstream changes in other vascular K^+^ currents or LTCCs.

### Differentiation of human iPSCs to vascular smooth muscle cells

To examine the electrophysiology of hiPSC-VSMCs, and to gain further insight to vascular consequences of CS variants, two genetically unrelated control human iPSC lines (C2a and AN-1.1 from individuals with no known disease-associated variants) were first differentiated to vascular smooth muscle cells. hiPSC-VSMCs were differentiated using the protocol developed by Patsch et al.^1^, which allows rapid and efficient differentiation to VSMCs that resemble native VSMCs based on expression of key marker genes, global transcriptomic and metabolomic signatures, and key functional characteristics including contractility and extracellular fibronectin deposition mediated by TGF-ß signaling^1^. This protocol is completely chemically defined and, in our hands, is reproducible under the published conditions. hiPSCs begin as relatively small, round, gray cells with dark nuclei (Fig. 2A). For four days, the cells are induced to become lateral mesodermal progenitors, which appear slightly enlarged but morphologically similar to hiPSCs. At this stage, the cells form a confluent monolayer, and patches of dead cells begin to accrue near the end of the 4-day lateral mesoderm induction. Finally, the cells are committed to a VSMC fate via exposure to PDGF-BB and Activin A. Once fully differentiated and cultured on a collagen-coated surface, the hiPSC-VSMCs possess a tapered, spindle-shaped morphology, and tend to align with one another, forming networks of multicellular whorls (Fig. 2B).

**Figure 2:**
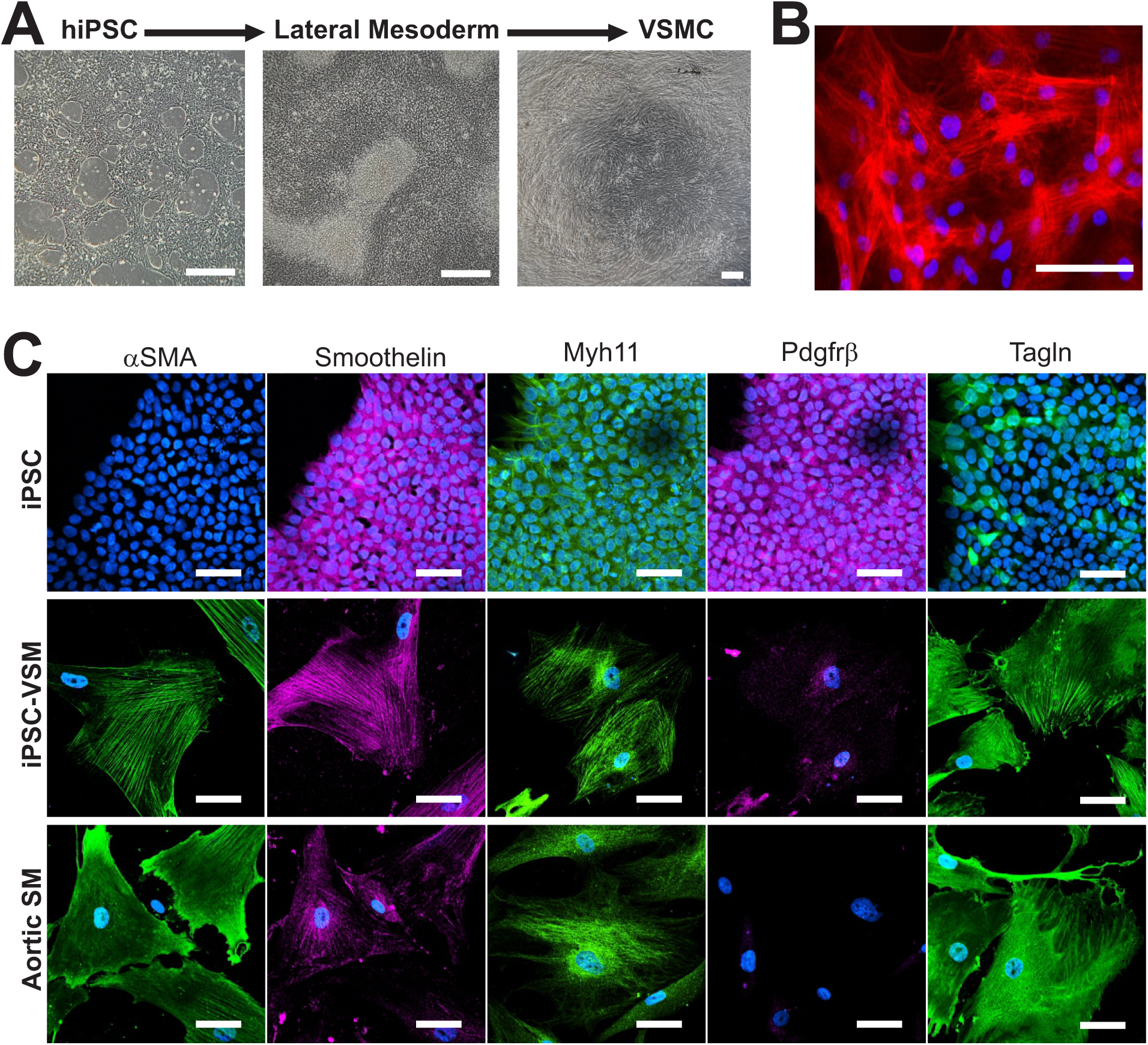
Differentiation of hiPSCs to VSMCs. (**A**) Images of (*left*) hiPSCs prior to differentiation, (*center*) after initial differentiation to lateral mesodermal progenitors, and (*right*) after full differentiation to hiPSC-VSMCs cultured on a collagen-coated surface (scale bars = 200 μm). (**B**) Composite fluorescent image of hiPSC-VSMCs with DAPI nuclear stain (blue) and Cy3-conjugated alpha-SMA stain (red), showing a dense network of discrete, co-aligned filaments expressed by hiPSC-VSMCs (scale bar = 50 μm). (**C**) Composite fluorescent images of undifferentiated iPSCs, hiPSC-VSMCs, and human aortic VSMCs, stained with DAPI nuclear stain (blue) versus antibodies for human αSMA (green), Smoothelin (magenta), Myh11 (green), Pdgfrβ (magenta), and Tagln (green) (scale bar = 50 μm in all panels).

We verified that the differentiation was effective by fixing undifferentiated iPSCs, iPSC-VSMCs, and cultured human aortic VSMCs, and staining for multiple VSMC markers: alpha-SMA (smooth muscle actin), Smoothelin, Myh11, Pdgfrβ, and Tagln. As shown in Fig. 2C, both the morphology and the VSMC protein expression pattern of human iPSC-VSMCs are strikingly similar to those of native human aortic VSMCs, with alpha-SMA filaments expressed as organized, co-aligned networks (Fig. 2).

### hiPSC-VSMCs express very similar ion channels to those in native mouse arterial VSMCs

To assess the vascular ensemble of K_v_ currents functionally expressed in hiPSC-VSMCs, recordings were obtained in these cells with the same protocols and solutions as above, in mouse VSMCs (Fig. 3). Consistently, we observed vascular K_v_ currents that were very similar to those measured in native VSMCs, with nearly identical current amplitudes and time-dependence (Fig. 3A). BK currents were also evident in some recordings, although less consistently than in mouse VSMCs, and were not characterized. We also detected nifedipine-sensitive LTCCs with essentially identical time dependence and current density to those in native mesenteric VSMCs (Fig. 3B).

**Figure 3:**
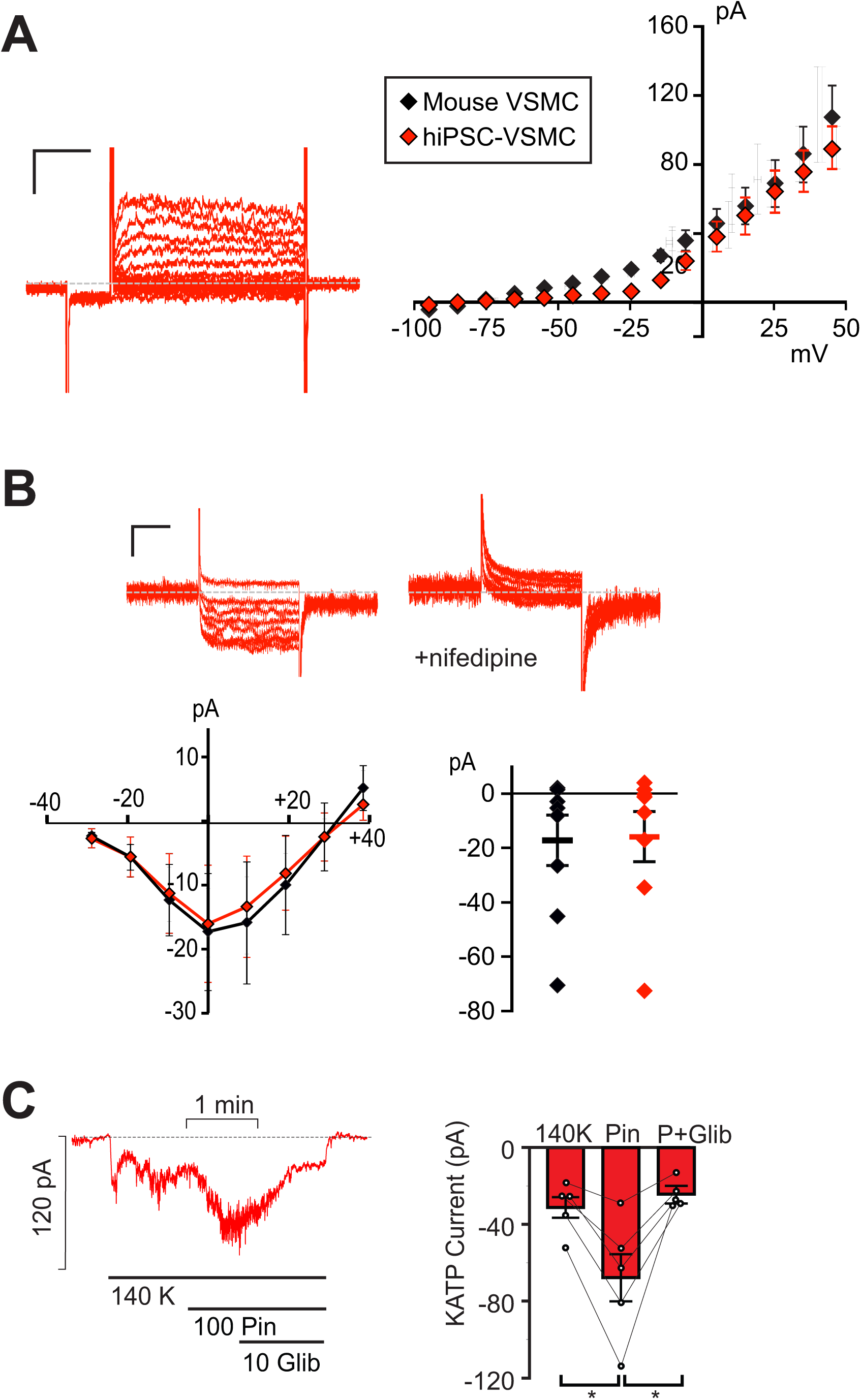
Electrophysiological characterization of hiPSC-VSMCs. **A**) (*left)* Representative whole-cell voltage-clamp recordings show K_v_ currents from a hiPSC-VSMC (scale bars = 50 pA, 100 ms). Voltage-clamp protocol was as in Figure 1. (*right*) Peak I-V relationship (measured during the final 100 msecs of the voltage pulse) shows mean ± s.e.m, for n = 9 cells, as well as the WT mouse I-V from Fig. 1. (**B**) (*top)* Representative whole-cell voltage-clamp recordings of LTCCs were obtained from a hiPSC-VSMC (scale bars = 10 pA, 75 ms). Voltage-clamp protocol as in Figure 2. Recordings were obtained in the presence of 1μM Bay K8644 (*left*), and conductance by LTCC was confirmed by >95% inhibition with 10μM nifedipine (*right*). (*bottom*) Peak I-V relationships in Bay K8644 are summarized (together with data from WT mouse VSMCs from Figure 2), with peak current at 0 mV displayed right for individual traces. Error bars show mean + s.e.m. (n = 8 WT and 6 VM cells in each case). (**C**) (*left*) Representative whole-cell voltage-clamp recordings of KATP channel conductance from control C2a hiPSC-VSMC using an intracellular pipette solution containing no nucleotides. Cells were voltage-clamped at - 70 mV. (*right*) Summary of mean currents in basal conditions, in 100μM pinacidil, and in 100μM pinacidil plus 10μM glibeclamide (mean + s.e.m., n = 5 cells). Statistical significance was determined by pairwise t-test with Bonferroni correction for multiple comparisons. * P<0.05, ** P<0.01, *** P<0.001.

These experiments provide a key first demonstration of hiPSC-VSMC electrophysiology, which shows similarity with native vascular ionic currents, essential for any future studies using these cells to understand vascular function in general, and particularly for those assessing vascular myocyte excitability, contractility, and response to electrically active agents.

### Pinacidil-sensitive K_ATP_ channels are present in hiPSC-VSMCs

Pinacidil-sensitive K_ATP_ channels, formed by Kir6.1 and SUR2 subunits, are key determinants of VSMC membrane voltage and excitability^28^. In previous studies, we characterized vascular K_ATP_ activity and pharmacology in mouse aortic VSMCs, with an intracellular pipette solution containing no ATP^18^. Using identical experimental conditions, we obtained whole-cell voltage-clamp recordings of basal K^+^ conductance in hiPSC-VSMCs (Fig. 4C). K^+^ current increased substantially when the SUR2-selective activator pinacidil (100μM) was applied, and current decreased following the addition of the K_ATP_ inhibitor glibenclamide (10μM) in the continued presence of pinacidil (Fig. 3C), reflecting activation of very similar K_ATP_ currents to those recorded in WT mouse VSMCs ^18^. Pinacidil responsivity indicates that hiPSC-VSMC K_ATP_ channels comprise the same vascular-type SUR2 isoform that is expressed in native VSMCs. Reduction of pinacidil-activated currents to essentially basal levels by glibenclamide further confirms that these currents are conducted by K_ATP_, with glibenclamide efficacy closely resembling that previously measured in native VSMCs^18^. Given these observations, hiPSC-VSMCs constitute a promising approach for the study of molecular and cellular consequences of CS variants.

**Figure 4:**
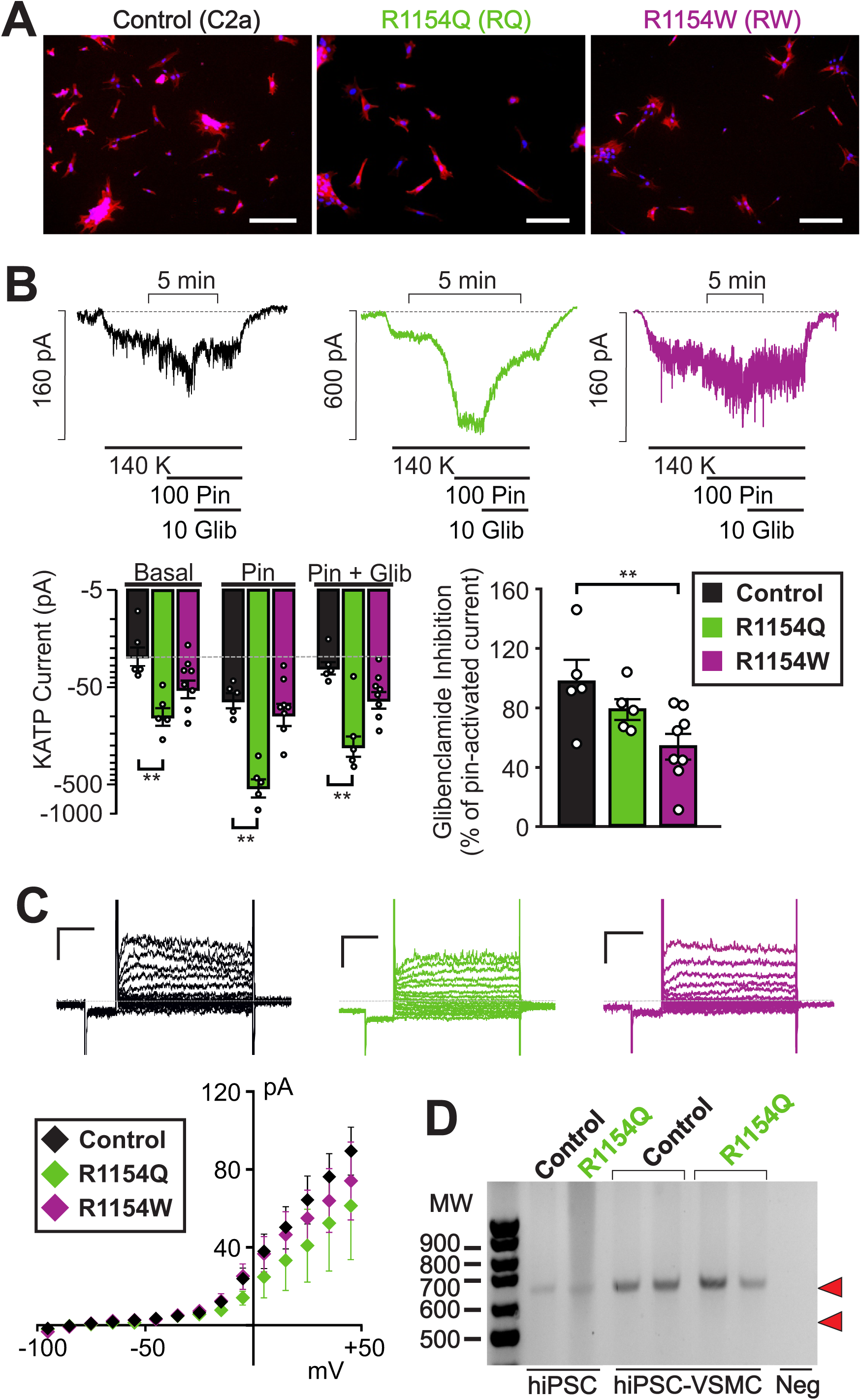
Electrophysiological characterization of CS hiPSC-VSMCs. (**A**) Composite fluorescent image of sparsely plated control C2a, R1154Q and R1154W hiPSC-VSMCs with DAPI nuclear stain (blue) and Cy3-conjugated alpha-SMA stain (red) (scale bar = 200 μm). (**B**) Representative whole-cell voltage-clamp recordings from control hiPSC-VSMCs (left), RQ hiPSC-VSMCs (middle), and RW hiPSC-VSMCs (right) using an intracellular pipette solution containing 100 μM MgATP and 500 μM MgADP. Cells were voltage-clamped at -70 mV. (*below*) Summary of mean currents in basal conditions, in 100μM pinacidil, and in 100μM pinacidil plus 10μM glibeclamide (mean + s.e.m., n = 5 cells in each case), as well as fraction of current inhibited by glibenclamide (*right*). (**C**) (*left)* Representative whole-cell voltage-clamp recordings show K_v_ currents from control, RQ and RW hiPSC-VSMCs (scale bars = 50 pA, 100 ms), voltage-clamp protocol as in Figure 1. (*below*) I-V relationships show mean ± s.e.m., for n = 9 (control), 5 (RQ) and 5 (RW) cells in each case. (**D**) cDNA PCR product from control and RQ hiPSCs reveals only a single band corresponding to a 642 bp fragment from full-length SUR2 cDNA (predicted site indicated by upper red arrowhead) and no band corresponding to the predicted 549 bp from exon 28-excluded cDNA (predicted site indicated by lower red arrowhead) (representative gel from 3 different differentiations of R1154Q patient hIPSC-derived VSMCs). For hiPSC-VSMCs, two control samples and two RQ samples are shown, generated from two separate differentiations. Statistical significance was determined by Mann-Whitney *U* test (α = 0.05). * P<0.05, ** P<0.01, *** P<0.001.

### Increased basal K_ATP_ activity, and decreased sensitivity to glibenclamide, in CS patient-derived hiPSC-VSMCs

We next developed human iPSC-derived vascular myocyte models for Cantú Syndrome using two patient-derived iPSC lines (which had been generated using Sendai virus–based reprogramming vectors on PBMCs and renal epithelial cells). These cells were obtained from CS patients carrying the R1154Q (RQ) and R1154W (RW) mutations, respectively, which are the first- and second-most common CS-associated variants^20^. Two subclonal hiPSC lines were produced for each mutation, and DNA sequencing analysis confirmed the expected mutation in each. Expression of human pluripotency-associated genes and a normal karyotype were confirmed for all hiPSCs prior to subsequent experiments, and CS hiPSC-VSMCs were generated as described above. Differentiation was highly efficient for both RQ and RW hiPSC-VSMCs, which were morphologically indistinguishable from control hiPSC-VSMCs, with alpha-SMA consistently expressed in control, RQ and RW hiPSC-VSMCs (>99% positive for each cell line, Fig. 4A).

To assess the molecular consequences of CS mutations in human iPSC-VSMCs, whole-cell voltage-clamp was used to measure K_ATP_ currents in RQ and RW hiPSC-VSMCs, as well as control C2a hiPSC-VSMCs (Fig. 4B). Experimental conditions were similar to those described above, but the pipette solution contained 100μM MgATP and 500μM MgADP (see Methods) to determine channel behaviors under nucleotide regulation, relevant to K_ATP_ activity *in vivo*. As shown in Fig. 4B, basal K^+^ conductances were elevated in each of the CS patient-derived hiPSC-VSMCs compared to control cells (>4-fold in RQ cells and >2-fold in RW cells, note logarithmic scale). Pinacidil-activated currents were ∼2.5x basal in WT (Fig. 4B) and ∼5.5x in RQ hiPSC-VSMCs (Fig. 4B), but only ∼1.9x in RW hiPSC-VSMCs (Fig. 4B).

In control hiPSC-VSMCs, glibenclamide inhibited almost 100% of the pinacidil-activated current, but glibenclamide action was reduced in mutant cells, inhibiting only ∼80% and ∼55% of pinacidil-activated current in RQ and RW, respectively (Fig. 4B). Increased basal currents in the CS variant cells are consistent with elevated basal K_ATP_ activity due to the known molecular consequences of the mutations^29, 30^. However, since glibenclamide fails to reduce these currents to basal levels, we cannot rule out the possibility that increased basal K^+^ currents are non-K_ATP_ mediated. To test this possibility, whole-cell voltage-clamp was used, as described above, to characterize the vascular ensemble of functional K_v_ currents with high (5mM) ATP in the pipette (Fig. 4C). I-V relationships revealed no significant basal K^+^ conductances. Voltage-activated K_v_ currents were non-significantly reduced in RQ, and RW hiPSC-VSMCs compared to control, confirming that elevated basal currents in Fig. 4B are indeed K_ATP_-mediated, and further suggesting that K_ATP_ GoF in CS vasculopathy does not lead to secondary, cell-autonomous changes in vascular K_v_ currents.

### Cell-autonomous consequences of genetic K_ATP_ overactivity underlie both hypomyotonic and hyperelastic components of CS vasculopathy

We previously generated a third CS mouse model in which the SUR2[R1154Q] was knocked-in to the endogenous locus^31^. In contrast to the mice carrying the SUR2[A478V] or Kir6.1[V65M] CS-associated variants, SUR2-dependent K_ATP_ currents and the resultant CS phenotype were both minimal in R1154Q mice. This was a result of alternate splicing leading to excision of exon 28, 3’ of the introduced variant, which led to non-functional SUR2^31^. If occurring in human patients with this variant, such splice excision could significantly ameliorate the CS features resulting from the molecular GoF itself. However, RT-PCR analysis of cDNA amplified from RNA isolated from control and RQ hiPSC-VSMCs (Fig. 4D) revealed only a single transcript band, with no evidence of exon 28 exclusion, in agreement with prior observations from primary human tissues samples^31^. This indicates that exon 28 exclusion is only seen in the R1154Q mouse and that a pure GoF phenotype will be present in human R1154Q cells. Consistent with this is a striking increase in K_ATP_ activity in RQ hiPSC-VSMC patch-clamp recordings (Fig. 4B), in contrast to native RQ mouse aortic VSMCs.

To understand how K_ATP_ GoF impacts VSMC excitability under physiological conditions, whole-cell current-clamp (*I*=0) was used to measure membrane voltage (V_m_) immediately (<2s) upon break-in. In the two genetically unrelated control C2a and AN-1.1 hiPSC-VSMC lines, we observed identical mean V_m_ values (-44 mV), consistent with reports of native VSMC resting V_m_, which lies in the range of -35 to -45mV^32–34^. Consistent with enhanced basal K_ATP_ conductance (Fig. 4B), both RQ and RW hiPSC-VSMCs were similarly hyperpolarized by 10-15mV relative to both controls (Fig. 5A).

**Figure 5:**
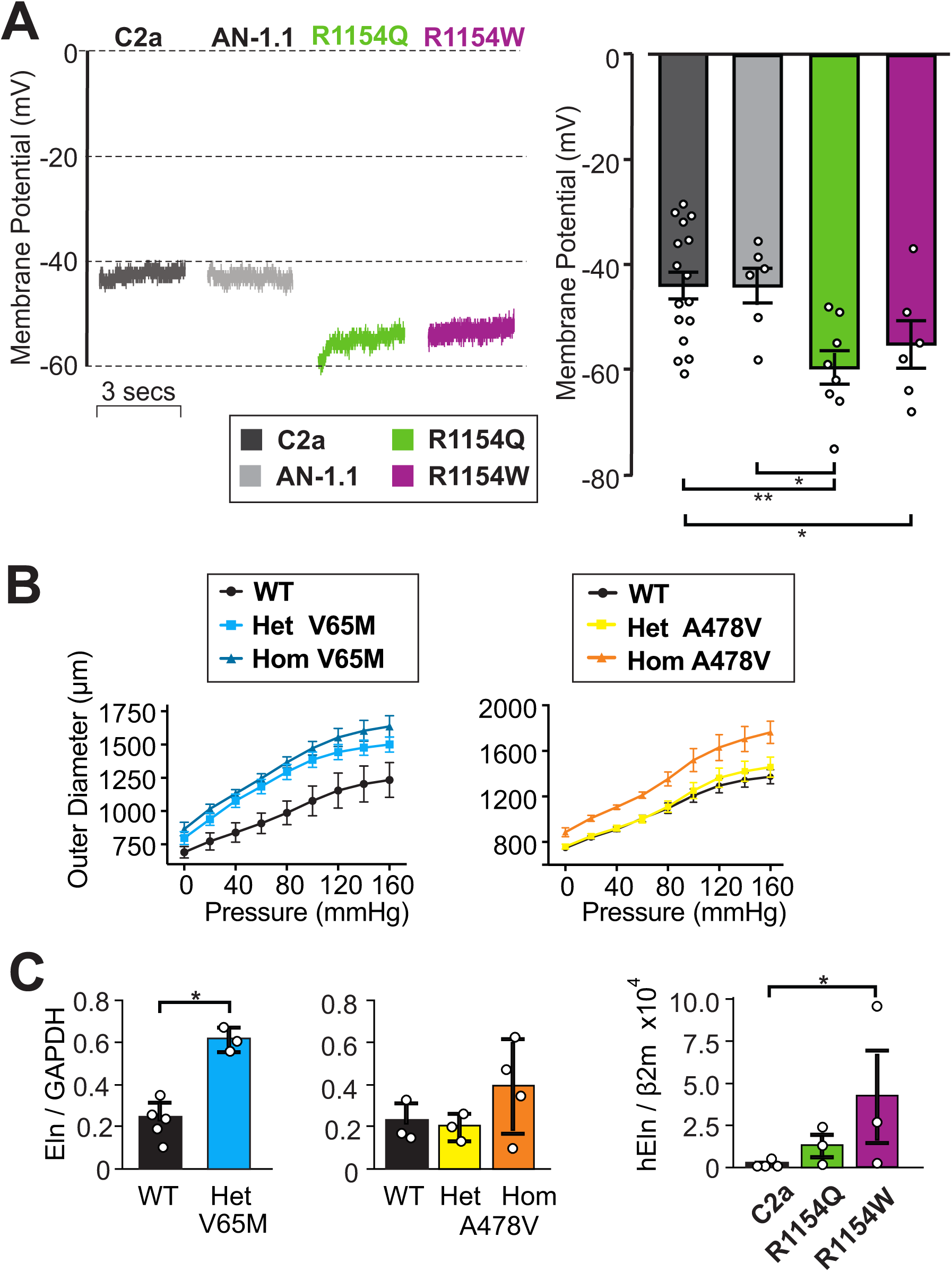
Cell-autonomous consequences of CS mutations on human iPSC-VSMC excitability and elastin expression. (**A**) (*above*) Representative whole cell current-clamp (at zero current) recordings from two control hiPSC-VSMC lines (*left*; C2a, AN-1.1) and two CS variant hiPSC-VSMC lines (*right*; RQ, RW) using an intracellular pipette solution absent of nucleotides. (*below*) Individual and mean (± s.e.m., n = 6-16 cells) initial Vm following break-in from experiments as above. (**B**) Vessel compliance in pressurized aortae of p21 WT, heterozygous and homozygous Kir6.1[V65M] mice (*left*), and WT, heterozygous and homozygous SUR2[A478V] (*right*) mice. Data shows mean + s.e.m., n= 4-5 animals in each case. (**C**) Elastin mRNA expression (normalized to GAPDH expression) in aortae from WT and V65M or A478V CS mice (left 2 panels), and from C2a control or R1154W and R1154Q hiPSC-VSMCs, normalized to β2 microglobulin (right panel). Statistical significance was determined by Mann-Whitney *U* test (α = 0.05). * P<0.05, ** P<0.01, *** P<0.001.

Finally, tortuous, dilated blood vessels with decreased PWV ^23, 24, 35^, as well as joint hyperelasticity^20, 36^ and tracheomalacia^20^, are major features of CS. Potentially, these might all be related to excess elastin production, which has been reported in response to treatment of blood vessels and cultured VSMCs with the K_ATP_ activators minoxidil^9–12^ and pinacidil^37^. As shown in Fig. 5B, compliance was increased in aortae from homozygous A478V and both hetero- and homozygous V65M mice at P21, the effect being proportional to the severity of the molecular consequences (from WT < Het < Hom in each case). In each case, the increased compliance was paralleled by proportionate trends in increased elastin mRNA expression in vessels at 3 months (Fig. 5C). We further measured mRNA expression from control (C2a), RQ and RW hiPSC-VSMCs. Elastin gene expression was barely detectable in control C2a samples, but it was increased in both of the CS lines, significantly so in RW (Fig. 5C). Taken together, these data are consistent with the increased elastin expression seen in CS being a direct, cell-autonomous, response to the increased VSMC K_ATP_ activity.

## Discussion

### The lack of electrical compensation for K_ATP_ GoF in CS

Blood pressure depends on vascular tone, which in turn is bidirectionally responsive to homeostatic needs, increasing or decreasing in response to even modest membrane depolarization or hyperpolarization from a potential (V_m_) that is typically close to the activation threshold for LTCCs^32–34^. In this voltage range, subtle changes in the activity of K_ATP_ or other channels, whether genetic or in response to neurohumoral signaling, will change vascular excitability, unless compensated by changes in another current. The finding that VSMCs from CS mice are basally hyperpolarized^31^, as are GI SMCs ^38^, is therefore consistent with enhanced basal K_ATP_ conductance in CS that is uncompensated by any change in either K_v_ currents or LTC currents. This is consistent with previous findings in cardiac muscle, which show no compensation for K_ATP_ subunit knockout by other K_ATP_ subunits, nor changes in K_v_ currents in response to K_ATP_ GoF in other K_ATP_ channel subunits^39, 40^. More broadly, genetic knockout of background Kir2.1 channels in vascular or cardiac myocytes are also not compensated for by changes in K_v_ or LTCCs^41, 42^.

Therefore, while we cannot completely rule out changes in all vascular ion channels, no changes in the major K_v_ and LTC currents that govern vascular excitability appear to contribute to, or compensate for, CS vasculopathy. Lack of compensation by other currents helps to explain why, despite the homeostatic imperative, manipulation of VSMC K_ATP_ conductance leads to marked changes in blood pressure in experimental animals, from a mean BP of ∼115 mM Hg in Kir6.1^-/-^ animals to ∼85 mmHg in smooth muscle-direct K_ATP_ GoF transgenic animals^43^. Interestingly, the results contrast with the finding that cardiac LTCCs are actually increased with cardiac K_ATP_ GoF^40, 44^, a potentially compensatory consequence in response to enhanced adrenergic signaling^44^.

### Human iPSC-derived VSMCs recapitulate native VSMC electrophysiology, resembling VSMCs from resistance vessels

A major goal of disease modeling in animals is to translate the findings to the human condition, and it is critical that the animal model be biologically comparable. Human VSMCs are not readily accessible, and the electrophysiology of human VSMCs has received relatively little attention^45–47^. Human iPSC-derived VSMCs therefore offer a potentially very profitable model in which to explore human VSMC biology in general, including electrophysiology. Although characterization of cellular electrophysiology is essential for use of these cells to model disease processes, there has been no study to date. We used an established protocol consisting of fully chemically defined conditions to differentiate hiPSCs into alpha-SMA-positive hiPSC-VSMCs via lateral mesodermal lineage. Subsequently, whole-cell patch-clamp electrophysiology was used to characterize functional expression of key channels underlying vascular excitability in identical conditions to those used for native mouse VSMCs discussed above. The ensemble of K_v_ currents in human iPSC-VSMCs closely resembles that in native mouse VSMCs, confirming that hiPSC-VSMCs recapitulate a key component of native vascular electrophysiology. Moreover, the non-inactivating time-dependent delayed rectifier K^+^ currents, as well as noisy, non-inactivating currents, which we ascribe to Ca^2+^-activated maxi-K (BK) channels, are quite similar to those originally reported in human mesenteric arterial myocytes^47^. In addition, LTCCs are detected at a similar levels, and with identical voltage-dependence to those in mouse mesenteric myocytes, and quite similar to those reported in human mesenteric arterial myocytes^48^. Having not detected LTCC in native aortic VSMCs, we suggest that increased functional LTCC expression may represent an electrical ‘signature’ of contractile VSMCs, and that hiPSC-VSMCs differentiated under the conditions used may more closely resemble VSMCs from resistance vessels than from elastic vessels. In their original report, Patsch et al.^1^ mentioned two potential options for hydrogel substrate: collagen, for promotion of a ‘contractile’ phenotype, and gelatin, for promotion of a ‘synthetic’ phenotype. In the present study, we cultured cells on collagen-coated surface, consistent with the presence of LTCCs, as in contractile myocytes.

### CS patient-derived hiPSC-VSMCs exhibit markedly elevated K_ATP_ activity, with decreased glibenclamide sensitivity

We also found that K_ATP_ channels are expressed in hiPSC-VSMCs at very similar levels to those in murine VSMCs^18^. Importantly, pharmacological sensitivity to the SUR2-selective drug pinacidil shows that these K_ATP_ channels comprise the appropriate ‘vascular-type’ architecture. In CS patient-derived hiPSC-VSMCs, K_ATP_ conductances were markedly elevated under basal conditions, more so in the RQ myocytes. This is consistent with the effect of both R1154W and R1154Q mutations to enhance Mg-nucleotide activation of recombinant K_ATP_ channels, particularly R1154Q^30^. Both mutations also increased the pinacidil-activated current – an effect that was much more prominent in RQ cells. Since pinacidil acts to stabilize Mg-nucleotide activated channels, this is also consistent with the much greater MgATP activating effect in R1154Q versus R1154W channels^30^. Finally, the relative inhibitory effect of glibenclamide was lower for the RQ, and particularly RW hiPSC-VSMCs, again consistent with findings from recombinant mutant channels^30^. This indicates that individuals with the two most common CS variants^20^, may receive limited therapeutic effect from glibenclamide and potentially other sulfonylurea drugs, and may instead benefit from drugs that act through a separate binding site and/or distinct allosteric mechanism.

### Distinct components of CS vasculopathy arise from cell-autonomous effects in VSMCs

Recent advances have made it increasingly clear that CS is primarily a smooth muscle-driven disease^18, 21, 49^, yet there has been a lack of fundamental understanding as to how CS mutations alter VSMC function. At least one component involves decreased SVR, which is a reflection of vascular tone, which in turn is determined by VSMC membrane potential. Importantly, control hiPSC-VSMCs from two genetically unrelated origins exhibit essentially the same membrane potential that is consistent with past reports in native VSMCs^32–34^, somewhat depolarized relative to that of non-vascular myocytes, and near the activation range for LTC channels. Consistent with K_ATP_ currents being major determinants of vascular membrane potential and hence excitability^50^, the resting membrane potential was markedly more negative in both CS lines, and can explain why vasodilation specifically, and smooth muscle function more generally, is a major driver of CS pathology^17, 20, 21 38^.

R1154Q and R1154W are the most common CS-associated variants in SUR2^20^ and cause very marked molecular GoF ^29, 30^. Unexpectedly, the phenotype of a CS mouse in which SUR2[R1154Q] was knocked in^31^ was minimal compared to mice carrying the CS-associated SUR2[A478V] or Kir6.1[V65M], due to alternate splicing leading to excision of exon 28, which led to non-functional channels. Such splice excision could significantly ameliorate the CS features resulting from the mutation itself if it happened in humans. However, the lack of any detectable exon 28 exclusion in hiPSC-VSMCs, or in CS patient biopsies^31^ suggests this is not occurring in humans.

CS vasculopathy includes not only the hypomyotonic component considered above, but also a hyperelastic component, characterized by tortuous vessels with increased compliance and decreased PWV, as well as aortic root dilation associated with aortic insufficiency, and aortic aneurysms ^23, 24, 35, 51, 52^. These latter phenomena are not trivially explained by an effect of K_ATP_ activation, but are consistent with multiple previous studies that have shown increased elastin expression in cultured cells and blood vessels exposed to pharmacological K_ATP_ activators, including minoxidil and pinacidil^9–12, 37^. Enhanced elastin mRNA expression, not only in mouse CS vessels, but also in CS variant hiPSC-VSMCs, implicates a cell-autonomous consequence of K_ATP_ GoF in VSMCs, independent of any systemic signaling. The connection between K_ATP_ activity and elastin gene expression remains unexplained, but potentially, like the hypomyotonic component, it is driven by effects of membrane potential on cellular calcium levels.

### Implications for hiPSC-VSMCs in the search for vascular therapeutics

Our results represent the first characterization of hiPSC-VSMC electrophysiology, and demonstrate that hiPSC-VSMCs recapitulate typical electrophysiology of native VSMCs, an important validation for the use of such cells for the study of human vascular pathologies in general. Our data also shed light on cell-autonomous phenomena that explain both the hypomyotonic and hyperelastic components of CS vasculopathy. While there is evidence that the hypomyotonic dysfunction can be treated with sulfonylurea inhibitors *in vivo*^53^, there is so far no evidence that the hyperelastic component can be corrected. Deficient elastin expression has been a key obstacle to the use of hiPSC-VSMCs for tissue-engineering^5^; our findings suggest that electrical excitability may be one potential avenue for increasing elastogenesis in hiPSC-VSMCs. Overall, our results provide a CS patient-derived human cell model suitable both for detailed study of the earliest mechanisms of wide-ranging CS cardiovascular pathologies, as well as for identification of new pharmacotherapeutic candidates.

## Novelty and Significance

### What is known?

- Cantu syndrome (CS), which includes low systemic vascular resistance (SVR) and increased vascular elasticity, is caused by gain-of-function in vascular K_ATP_ channels
- The K_ATP_ channel minoxidil causes increased elastin production in vascular smooth muscle (VSM)

### What new information does this article contribute?

- Major vascular ion channels are unaltered in CS mouse vascular smooth muscle cells (VSMCs) by CS
- This first study of human iPSC-VSMC electrophysiology reveals expression of essentially the same excitable currents as primary murine VSMCS, including K_v_, LTCC, and K_ATP_, as well as similar membrane voltage (V_m_)
- Human CS patient-derived iPSC-VSMCs show basal K_ATP_ activation, but other major ion channels are unaffected, resulting in cell-autonomous hyperpolarization, explaining low SVR
- CS mouse VSMCs and human CS patient-derived iPSC-VSMCs both show increased elastin production (homozygous > heterozygous), showing that this is a cell-autonomous response to K_ATP_ activation

## Novelty and Significance

Cantu Syndrome (CS), which is caused by gain-of-function in vascular K_ATP_ channels, is characterized by large-vessel dilatation with decreased pulse-wave velocity, as well as low systemic vascular resistance, reflecting hyperelastic and hypomyotonic components. First, we have shown that the functional expression of voltage-gated K^+^ and Ca^2+^ currents are essentially the same in mouse vascular smooth muscle cells (VSMCs) from WT and CS mice, revealing lack of any compensatory electrical response. We then derived VSMCs from human induced pluripotent stem cells (hiPSC-VSMCs) and made a first characterization of their electrophysiology (EP), showing that functional expression of voltage-gated K^+^ and Ca^2+^ currents in control hiPSC-VSMCs is very similar to that in mouse arterial VSMCs. Both basal and pinacidil-activated K_ATP_ currents were considerably larger in CS hiPSC-VSMCs. Consistent with lack of cell-autonomous modulation of other currents, this resulted in membrane hyperpolarization, explaining the hypomyotonic basis of CS vasculopathy. Consistent with the hyperelastic component of CS, we also observed increased compliance and dilatation in isolated CS mouse aortas, which was associated with increased elastin mRNA expression (homozygous > heterozygous). We also found increased elastin mRNA in two genetically unrelated lines of CS hiPSC-VSMCs. These results show that increased VSMC elastin expression is also a cell-autonomous consequence of K_ATP_ channel activation.

## Author contributions

AMH, CMcC, CGN originally conceived the study; DKG was responsible for clinical contact with patients; AMH, K-CW, JRS developed the iPSC-VSMCs; AMH carried out all electrophysiology; AMH, SC and ANS carried out histology; AMH, CGN carried out analysis; CMH carried out compliance measurements; AMH, CMH carried out expression analysis; AMH, CGN wrote the paper, which was edited by all authors.

## Acknowledgements

This work was supported by NIH grants R35 HL140024 to CGN and R21 HD103347 to CGN/DKG, and by Pilot and Feasibility Grant (CIMED-18-04) to CH. AH was supported by NIH T32 HL125241. CMcC was supported by NIH grant K99 HL150277. CMH was supported by NIH K08 HL135400. We are very grateful to Kristina Hinman and Robert Mecham (Department of Cell Biology and Physiology) for help with compliance measurements.

